# On the optimal response vigor and choice under variable motivational drives

**DOI:** 10.1101/057208

**Authors:** Amir Dezfouli

## Abstract

Within a rational framework, a decision-maker selects actions based on the reward-maximisation principle, i.e., acquiring the highest amount of reward with the lowest cost. Action selection can be divided into two dimensions: (i) selecting an action among several alternatives, and (ii) choosing the response vigor, i.e., how fast the selected action should be executed. Previous works have addressed the computational substrates of such a selection process under the assumption that outcome values are stationary and do not change during the course of a session. This assumption does not hold when the motivational drive of the decision-maker is variable, because it leads to changes in the values of the outcomes, e.g., satiety decreases the value of the outcome. Here, we utilize an optimal control framework and derive the optimal choice and response vigor under different experimental conditions. The results imply that, in contrast to previous suggestions, even under conditions that the values of the outcomes are changing during the session, the optimal response rate in an instrumental conditioning experiment is a constant response rate rather than decreasing. Furthermore, we prove that the uncertainty of the decision-maker about the duration of the session explains the commonly observed decrease in response rates within a session. We also show that when the environment consists of multiple outcomes, the model explains probability matching as well as maximisation choice strategies. These results, therefore, provide a quantitative analysis of optimal choice and response vigor under variable motivational drive, and provide predictions for future testing.

## Introduction

According to the normative theories of decision-making, actions made by humans and animals are chosen with the aim of earning the maximum amount of future rewards while incurring the lowest cost (Neumann & Morgenstern, 1947). Within such theories, individuals optimize their actions by learning about their surrounding environment and in order to satisfy their long-term objectives. Such a problem, i.e., finding the optimal actions, is argued to have two aspects: (1) choice, i.e., which action among several alternatives should be selected, (2) response vigor, i.e., how fast the selected action should be executed. For example for a rat in a Skinner box with two levers, where pressing each lever delivers a reward with a certain probability, the problem of finding the optimal actions involves selecting a lever (choice) and deciding about the response rate on the chosen lever (response vigor). High response rates can have high costs (e.g., in terms of energy consumption), while on the other hand a low response rate implies an opportunity cost since the experiment session may end before the animal has earned enough reward. Optimal actions provide the right balance between these two factors, and based on the reinforcement-learning (RL) framework and methods from optimal control theory, characteristics of the optimal actions and their consistency with the experimental studies have been elaborated in previous works (Dayan, 2012; Niv, Daw, Joel, & Dayan, 2007; Salimpour & Shadmehr, 2014).

A major assumption in the previous models is that the values of the outcomes are stationary, and they do not change on-line in the course of a decision-making session. To see the limitations of such an assumption, imagine the rat is in a Skinner box and has started to earn outcomes (e.g., food pellets) by taking actions. One can assume that as a result of consuming rewards the motivation of the animal for earning more outcomes will decrease (e.g., because of satiety effects) and, therefore, the outcomes earned will have a lower value, which can potentially affect the optimal choice and vigor (Killeen, 1995b). Such effects, however, are not incorporated in previous models of response vigor, and it is assumed that the value of the outcomes is constant during the course of the decision-making session. On the other hand, the effect of such motivational changes on the choice between actions has been addressed in some previous models (Keramati, 2011; Keramati & Gutkin, 2014), however, their role in determining the response vigor has not yet been investigated. Here, building upon the previous works, we formulate the problem of response vigor and choice under changing motivational drives in an optimal control framework, and we derive the optimal response vigor and choice strategy under different conditions. We will show that even when the motivational drives are changing, the optimal response rate in an instrumental conditioning experiment is a constant response rate, and then we will elaborate how this prediction can be reconciled with the experimental data. The optimal predictions under different choice situations is also explored, and their relation to the empirical evidence is investigated.

## The Model

### The reward

It is assumed that at each point in time the decision-maker has a position in the outcome space denoted by *x_t_*, which represents the amount of outcomes gained up to time *t*. For example, if the outcome is water, *x*_*t*_ = 1 indicates that one unit of water has been gained up to time *t*. For simplicity, we assume that only one outcome is available, and thus the outcome space is one dimensional. In the next sections the model will be extended to the environments with multiple outcomes. The rate of outcome earning is denoted by *v*_*t*_, which represents how fast the decision-maker is moving in the outcome space (i.e., the velocity in the outcome space); for example if a rat is earning one unit of an outcome per unit of time then *v*_*t*_ = 1. Furthermore, we assume that there exists a *reward field,* denoted by *A_x,t_*, which represents the per unit value of the outcome for the decision-maker. For example if the outcome is food pellets, then *A_x,t_* represents the value of one unit of the food pellet at time *t*, given that the decision-maker has already earned *x* units of food pellets. As such, *A_x,t_* is a function of both time and the amount of outcome earned. This represents the fact that (i) the reward can change as a result of consuming previous outcomes (dependency on *x*); for example the reward of food pellets can decrease as an animal consumes more outcomes and becomes sated, and (ii) the reward of an outcome can change purely by the passage of time, as for example an animal can get more hungry by the passage of time, and therefore the reward of food pellets will increase (dependency on *t*).

In general, we assume that *A_x,t_* has two properties:

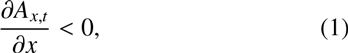

which entails that the values of the outcomes decrease as more outcomes are earned. Secondly, we assume that:

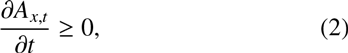

which entails that the values of outcomes do not decrease with the passage of time. This term, for instance, corresponds to the metabolic rate of the decision-maker. Given the reward field *A*_*x,t*_, the reward of gaining *δx* units of outcome will be *A*_*x,t*_*δx.*

The dependency of the reward field on the amount of outcome earned is indirect and it is through the motivational drive. In a computational model based on **RL** framework, Keramati and Gutkin (2014) provided a quantitative link between reward and motivation. In line with Hull (1943), they conceptualized the motivational drive as the deviations of the *internal states* of a decision-maker from their homeostatic set-points. For example, let’s assume that there is only one internal state, say hunger, where *H* denotes its homeostatic set-point, and there is an outcome which consuming each unit of it satisfies *l* units of the internal state. In this condition, the motivational drive at point *x*, denoted by *D_x_,* will be:

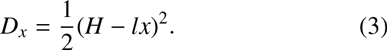

Keramati and Gutkin (2014) showed that such a definition of the motivational drive has implications that are consistent with the behavioral evidence. According to the framework, the reward generated by earning *δx* units of the outcome is proportional to the change in the motivational drive, which can be expressed as:

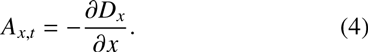

The above equation will be used for linking the consumption of outcomes to the changes in the motivational drive and the reward field.

### The cost

Using RL framework, previous studies have derived the optimal choice and response vigor in instrumental conditioning experiments, in which an animal is required to press a lever to earn rewards (Dayan, 2012; Niv et al., 2007). The optimal rate is argued to be the result of a trade-off between two factors: the cost of lever presses, and the opportunity cost of a certain response rate. The cost component is expressed as a function of the delay between consecutive lever presses. That is, if the previous lever press has occurred *τ* time steps ago, then the cost of the current lever press will be:

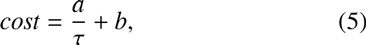

where *a* and *b* are constants. *b* is the constant cost of each lever press, which is independent of the delay between lever presses. The factor *a*, on the other hand, controls the rate-dependent component of the cost. According to the above relation, the higher the delay between lever presses the lower the cost will be. However, a long delay has an opportunity cost, i.e., by waiting a long time the animal is missing an opportunity to earn rewards that potentially could be obtained during the delay between presses. The optimal rate, therefore, will be determined by a trade-off between the cost of a certain response rate and its opportunity cost.

In the above framework, the target of the decision-making process is to find the optimal value of *τ*. Here, we express the cost as a function of the rate of outcome earning instead of the rate of action executions. We denote the cost function with *K_v_*, which indicates the cost of earning one unit of the outcome at rate *v*. In the case of a fixed-ratio (FR) schedule of reinforcement, in which the decision-maker is required to perform exactly *k* responses in order to earn an outcome, or in the case of a random-ratio (RR) or a variable ratio (VR) schedule, in which the decision-maker is required to perform on average *k* responses to earn an outcome, the cost defined in equation 5 will be equivalent to:

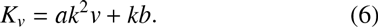

See Appendix A for the proof. In general, we assume that *Kv* satisfies the following relation:

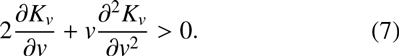

The above relation is required for deriving the results provided in the next sections. For example in the case of the cost function defined in equation 6, the above relation requires that *ak^2^* > 0, which implies that in the experiment at least one response should be required to earn an outcome (*k* > 0), and the cost of earning outcomes should be non-zero (*a* > 0).

Given *K_v_*, the cost of gaining *δx* units of outcome will be*K_v_δx*.

### The value

Based on the definition of the reward and the cost, the net amount of reward earned at each time step will be the reward earned (*vA_x,t_*) minus its cost (*vK_v_*). We denote this quantity by *L*:

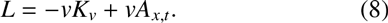

Now let’s assume that a decision-maker starts at point xo in the outcome space, and the total duration of the experiment session is *T*. We denote the total reward gained in this period with *S* _0,*T*_, which is the sum of the net rewards earned at each point in time:

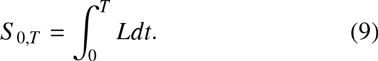

The quantity *S* _0,*T*_ is called the *value* function, and the goal of the decision-maker is to find the optimal rate of outcome earning that yields the highest value (*S* _0,*T*_). The optimal rates can be found using different variational calculus methods such as the Eular-Lagrange equation, or the Hamilton-Jacobi-Bellman equation (Liberzon, 2011). Here we use the Eular-Lagrange form, which sets a necessary condition for *δS* = 0:

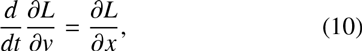

which implies that:

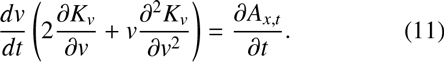

See Appendix A for the proof. Furthermore, since the endpoint of the trajectory is free (the amount of outcomes that can be gained during a session is not limited, but the duration of the session is limited to *T*), the optimal trajectory will also satisfy the transversality conditions:

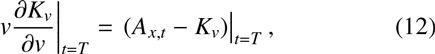

where as mentioned *T* is the total session duration. We will use equations 11,12 in order to derive the optimal response rates in different conditions.

## Optimal response vigor

In this section, we analyze the optimal response vigor in the condition that there is only one outcome and one response available in the environment, i.e., the outcome space has only one dimension. The analysis is divided into two sections. In the first section, the decision-maker assumes that the duration of the session is fixed, which will be extended in the next section to the conditions that the decision-maker assumes a probabilistic distribution over the session length.

### Fixed session duration

The optimal rate of outcome earning should satisfy equation 11. In the equation, the term ∂*A_x,t_*/∂*t* is the time-dependent change in the value of the outcome, and the term *dv/dt* is the rate of change in the rate of outcome earning. In the particular case that the time-dependent change in the outcome reward is negligible ∂*A_x,t_*/∂*t* = 0, then the only valid solution to equation 11 is *dv/dt* = 0 (see Appendix A). Thus, the optimal rate of outcome earning (and the optimal response rate) is constant throughout the session. This relation holds even in the condition that the reward of the outcome decreases throughout the session as a result of earning outcomes, e.g., because of the satiety effect. The following theorem summarizes this result:

#### Theorem 1

*If the reward field and the cost function satisfy equations 2 and 7, then the optimal rate of outcome earning will be non-decreasing* (*dv*/*dt* ≥ 0). *In the special case that the time-dependent change in the reward field is zero* (∂*A_x,t_*/∂*t* = 0), *then the optimal rate of outcome earning is a constant rate* (*dv*/*dt* = 0). *Furthermore, the optimal rate v*\* *satisfies the following equation:*

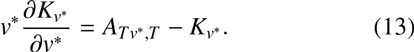

See Appendix A for the proof. As an intuitive explanation for why the constant rate is optimal, imagine a decision-maker who chooses a non-constant rate of outcome earning, and it earns in total *x_T_* amount of outcome during the session. If instead of the non-constant rate, the decision-maker selects the constant rate *v* = *x_T_/T*, then the amount of outcome earned will be the same as before; however, the cost paid during the session will be lower because the cost is a quadratic function of the outcome rate, and therefore, it is better to earn outcomes at a constant rate.

As an example, let’s assume that there is one internal state, say hungry, and the motivational drive, and the cost function are the ones defined in equations 3 and 6. Using Theorem 1 the optimal rate of outcome earning will be^1^:

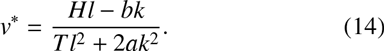

The response rate, in the case of the FR schedule can be obtained by multiplying the outcome rate by a factor of *k*, the number of responses required to earn an outcome:

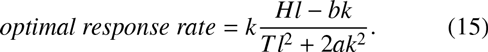

Equation 14 and 15 layout a quantitative relation between various parameters, and the optimal rate of outcome earning (equation 14), and the optimal rate of responding (equation 15), which can be compared against experimental evidence. Unfortunately, experimental data on the effects of different parameters on response rates are not consistent across studies, making it hard to compare the optimal actions with empirical data. Besides that, there is a further complexity due to the inconsistency in how different experiments have calculated response rates. In general, in an instrumental conditioning experiment, the duration of the session can be divided into three sections: (i) outcome handling/consumption time, which refers to the time that an animal spends on consuming the outcome, (ii) *post-reinforcer pause*, which refers to a pause after consuming the outcome and before starting to make responses (e.g., lever presses). Such a pause is consistently reported in previous studies using the FR schedule, (iii) *run time*, which refers to the time spent on making responses (e..g, lever presses). Experimental manipulations have been shown to have different effects on the duration of these three sections of the session, and whether each of these sections are included when calculating the response rates can affect the measurement of the results. In the following sections, we briefly review the currently available data from instrumental conditioning experiments and their relation with the predictions of the model.

**The effect of response cost** (*a* **and** *b*). Experimental studies in rats in a FR schedule (Alling & Poling, 1995), indicate that increasing the force required to make responses causes increases in both inter-response time and post-reinforcement pause. The same trend has been reported in Squirrel monkey (Adair & Wright, 1976). Consistently, the present model implies that increases in the cost of responses, for example by increasing the effort required to press the lever (increases in *a* and *b*), lead to a lower rate of outcome earning and a lower rate of responses (Figure 1).

**The effect of ratio-requirement** (*k*). Experimental studies mainly imply that the rate of outcome earning decreases with increases in the ratio-requirement (Aberman & Salamone, 1999; Barofsky & Hurwitz, 1968), which is consistent with the general trend of the optimal rate of outcome earning implied by the present model (as suggested by equation 14).

Experimental studies on the rate of responses, in the FR schedule, indicate that the post-reinforcer pause increases with increases in the ratio-requirement (Ferster & Skinner, 1957, Figure 23)(Felton & Lyon, 1966; Powell, 1968; Premack, Schaeffer, & Hundt, 1964). In terms of the overall response rates, some experiments report that response rates increase with increases in the ratio-requirement up to a point and beyond that point the response rates will start to decrease, in rats (Barofsky & Hurwitz, 1968; Kelsey & Allison, 1976; Mazur, 1982), pigeons (Baum, 1993) and mice (Greenwood, Quartermain, Johnson, Cruce,& Hirsch, 1974), although other studies have reported inconsistent results in pigeons (Powell, 1968), or a decreasing trend of response rates with increases in the ratio-requirement (Felton & Lyon, 1966; Foster, Blackman, & Temple, 1997). The inconsistency is partly due to the way that the response rates are calculated in different studies; for example in some studies the outcome handling and the consumption time are not excluded when calculating response rates (Barofsky & Hurwitz, 1968), in contrast to the other studies (Foster et al., 1997). As such any implication of the experimental data about the relationship between response rates and the ratio-requirement is not conclusive.

**Figure 1.**
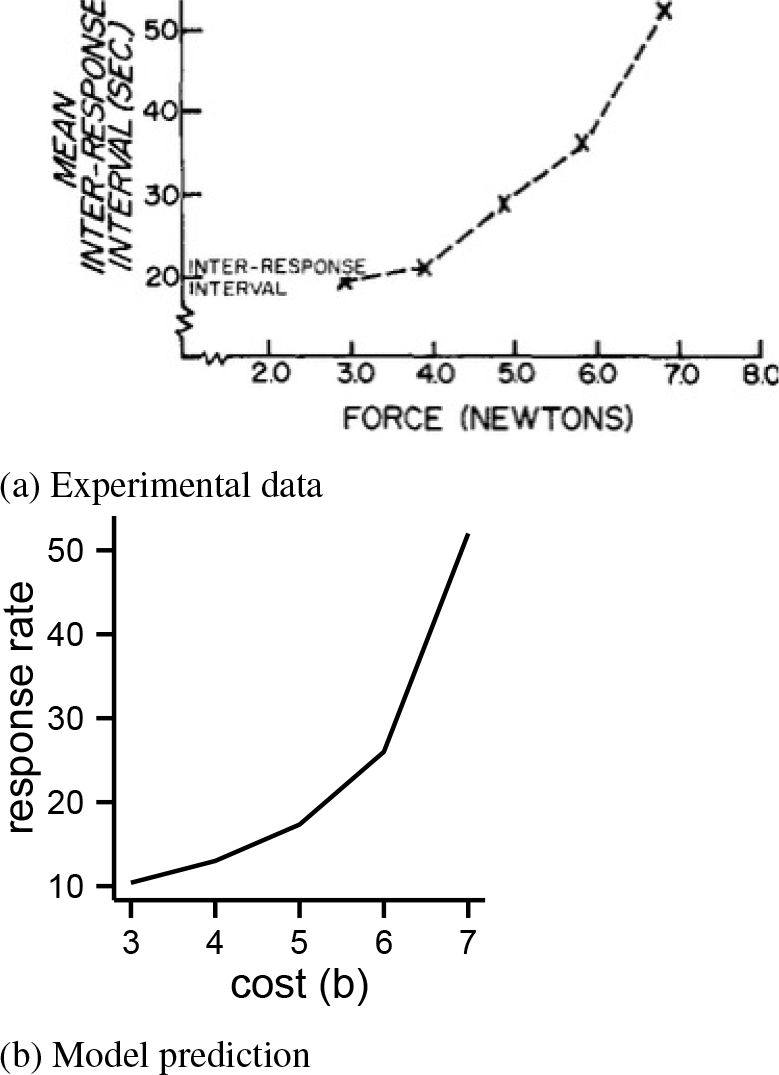
Effect of response cost on response rates. (a) Empirical data. Inter-response intervals when the force required to make a reponse is manipulated. Figure is adopted from Adair and Wright (1976). (b) Model prediction. Interresponse interval (equal to the inverse of response rates) as a function of cost of responses (b). Parameter values used for the simulation are *T* = 50, *k* = 1, *l* = 1, *a* = 1, *H* = 8.

With regard to the present model, it predicts the relationship between response rates and the ratio-requirement is an inverted U-shaped pattern (Figure 2a), consistent with some of the studies mentioned before. The exact relationship between the two factors depends on the value of the parameters. Generally speaking, if the rate-dependent cost of the responses is negligible (*a* = 0), the response rates will peak at *k* = *Hl* = (2*b*) and will start to decrease after that. As such, the prediction of the model can be consistent with the experiments mentioned before, depending on the initial motivational drive (*H*), the constant cost of responses (*b*), and the range of tested ratio-requirements.

**Figure 2.**
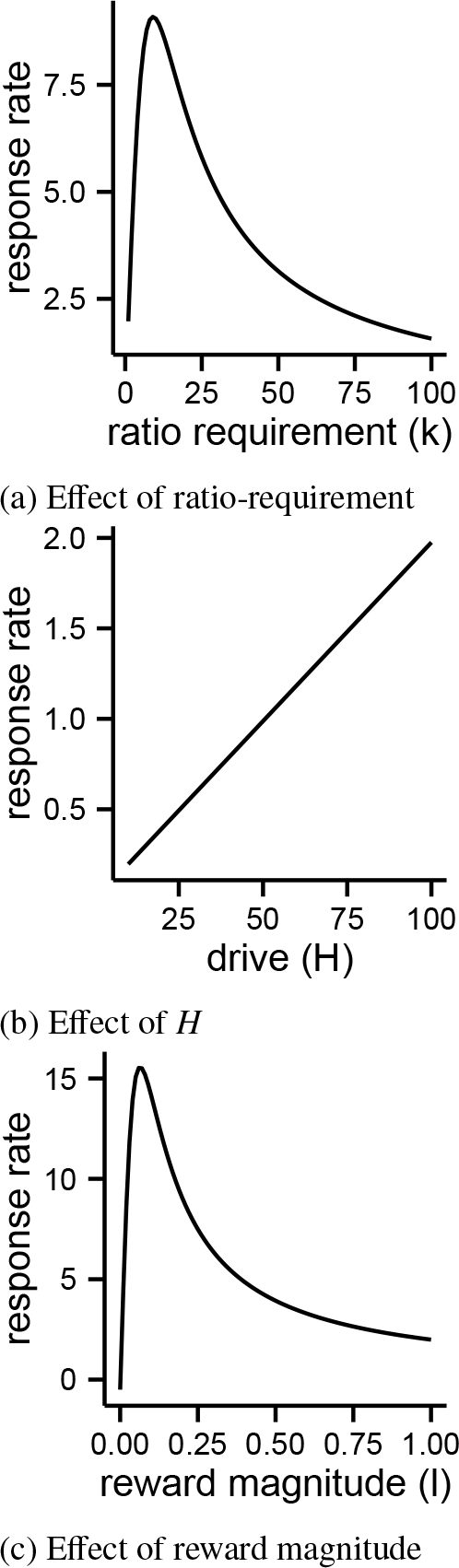
(a) The effect of ratio-requirement on the response rates. Parameters used for the simulations are *T* = 50, *l* = 1, *a* = 0:3, *b* = 0:05, *H* = 100. (b) The effect of the initial motivational drive on response rates. Parameters used are *T* = 50, *k* = 1, *l* = 1, *a* = 0:3, *b* = 0:05. (c) The effect of the reward magnitude on response rates. Parameters used are *T* = 50, *k* = 1, *a* = 0:1, *b* = 0:1, *H* = 100.

**The Effect of initial motivational drive** (*H*). Experimental studies in the case of a FR schedule suggest that response rates increase with increases in deprivation levels (Ferster & Skinner, 1957, Chapter 4)(Sidman & Stebbins, 1954). However, such increases are mainly due to decreases in the post-reinforcement pauses, and not due to the increases in actual rate of responses after the pause (see (Pear, 2001, Page 289) for a review of other reinforcer schedules). Consistently, the model predicts that with increases in the initial motivational drive (*H*), the rate of responses and outcome earning will increase (Figure 2b).

**The effect of reward magnitude** (*l*). Some studies show that post-reinforcement pauses increase as the magnitude of the reward increases (Powell, 1969), while other studies conclude that the post-reinforcement pause decreases (Lowe, Davey, & Harzem, 1974), although in the later study the magnitude of the reward is manipulated within a session and a follow-up study showed that at the steady state, the post-reinforcement pause increases with increases in the magnitude of the reward (Meunier & Starratt, 1979). The reward magnitude, however, does not have a reliable effect on the overall response rates (Keesey & Kling, 1961; Lowe et al., 1974; Powell, 1969). Regarding the prediction of the model, the effect of the reward magnitude on the outcome and response rates is an inverted U-shaped relationship (Figure 2c), with the peak at *l* = 2*bk*/*H* (assuming *a* = 0), and therefore depending on the value of the parameters the predictions of the model can be consistent with the experimental data.

**Within session pattern of responses**. It has been established that in various schedules of reinforcement (McSweeney & Hinson, 1992; McSweeney, Weatherly, & Swindell, 1995), including the VR schedule (McSweeney, Roll, & Weatherly, 1994), the rate of responses within a session has a particular pattern: the response rate reaches its maximum within a short delay after the session starts (warmup period), and then it gradually decreases toward the end of the session (Figure 3a). Killeen (1994) proposed a mathematical description of this phenomenon, which can be expressed as follows (Killeen & Sitomer, 2003):

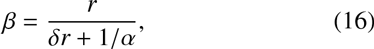

where *β* is the response rate, *δ* is the minimum delay between responses, *r* is the rate of outcome earning, and α is called *specific activation*^2^. The model suggests that as the decision-maker earns outcomes during the session, the value of *α* gradually decreases because of the satiety effect, which will cause decreases in response rates. Here satiety refers to both post-ingestive factors (such as blood glucose level; Killeen (1995a)) and/or pre-ingestive factors (for example sensory specific satiety; McSweeney (2004)). This model has been shown to provide a quantitative match to the experimental data. This explanation, however, is not consistent with Theorem 1, which posits that even in the condition that the motivational drive is changing within a session the optimal response rate is non-decreasing throughout the session, and, therefore, according to the present model the cause underlying the decreases in within-session response rates cannot be purely the changes in the motivational drive.

**figure 3.**
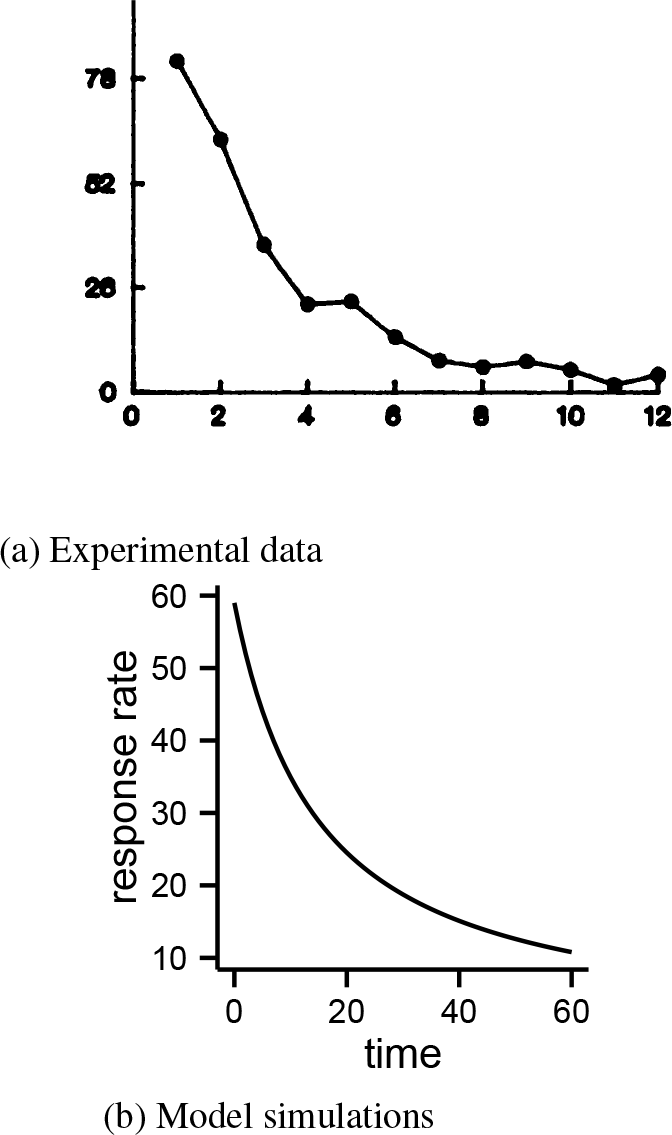
The pattern of within-session response rates. (a) Experimental data. The rate of responding per minute during successive intervals (each interval is 5 minutes) in a variable ratio (VR) schedule. The figure is adopted from McSweeney et al. (1994). (b) Theoretical pattern of within-session responses predicted by the model. Parameters used for the simulations are *k* = 15, *l* = 0.1, *a* = 0.002, *b* = 0.1, *H* = 900.

The optimal response rates provided by Theorem 1 are not consistent with the behavioral observations showing that the rate of responses for earning an outcome in actual fact decrease, and therefore, there is a clear inconsistency between predictions of the model and empirical data. Based on this, some of the assumptions made to develop the model should be violated by the decision-maker. The form of the cost and, reward functions is reasonably general. However, we assumed that the duration of the session, *T*, is fixed and known by the decision-maker. In the next section, we show that relaxing this assumption to the condition that the decision-maker is uncertain about the duration of the session will lead to predictions similar to the experimental data.

### Uncertain session duration

In the previous sections we considered conditions in which the duration of the session was fixed and known by the decision-maker. In this section we focus on the conditions that the decision-maker is uncertain about the session duration, i.e., the session duration follows a probability distribution function, that we denote by *p*(*T*). Under this assumption, the value of a trajectory starting at time *t* = 0 will be:

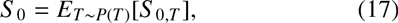

where *E* is the expectation. The value, therefore, is the average of the value of all the trajectories starting from point *t* = 0 and ending at time *t* = *T*, for all *T* > 0, weighted by the probability that the session will end at time *T*. The optimal trajectory that maximizes the above value is presented in Appendix A. However, here we use a more intuitive approximation of the above value function:

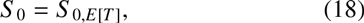

which implies that the decision-maker calculates the values based on the assumption that the session will end at *t* = *E*[*T*], i.e., it sets an expectation on how long the session will last and calculates the optimal response rates based on that expectation. Based on this, if *t*’ time step has passed since the beginning of the session, the value of the rest of the session
will be as follows:

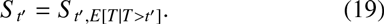

The following theorem maintains that the optimal rate of outcome earning that maximizes *S* _*t*’_ is a decreasing function of *t*’:

#### Theorem 2

*Given the following value function:*

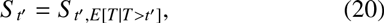

*and assuming that (i) the reward field and cost function satisfy equation 2 and 7 (ii) the time dependent change in the reward field is zero* (∂*A_x,t_*/∂*t* = 0), *and (iii) the probability that the session ends at each point in time is non-zero* (*p*(*T*) > 0), *then the optimal rate of outcome earning is a decreasing function of t*’:

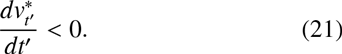

See Appendix A for the proof. As an intuitive explanation, it is apparent from equation 14 that the optimal rate of outcome earning is a decreasing function of the session length, i.e., when the session duration is long the decision-maker can take its time and earn outcomes more slowly. On the other hand, when the decision-maker is uncertain about the session duration, as time passes within the session the decision-maker’s expectation of the session duration increases. This phenomenon, which has been previously elaborated within
the context of delay gratification (McGuire & Kable, 2013; Rachlin, 2000), is more significant if the decision-maker assumes a heavy-tail distribution over *T*. In this condition as the time passes, the decision-maker will expect the session to last longer. This property, in addition to the fact that the optimal response rate is a decreasing function of the session duration, entails that as the time passes the optimal response rate will decrease. Based on this explanation, what underlies decrements in response rates within a session, in a normative perspective, is in fact the uncertainty in the duration of the session, and not the satiety effect.

For the simulation of the model, following McGuire and Kable (2013), we characterized the session duration using a Generalized Pareto distribution. Simulations of the model are depicted in Figure 3b, which shows that as time passes the optimal rate of responses decreases, consistent with the experimental data (see Appendix A for details).

## Optimal choice and response vigor

In the previous sections we assumed that the environment contained only one outcome, and the decision-maker only needed to decide about the response rates along one dimension. In this section we assume that there are multiple outcomes available in the environment, and the decision-maker needs to make decisions about the response rates along each outcome dimension. The position of the decision-maker instead of being a scalar will be a vector, denoted by **x**,which represents the amount of outcomes earned along each outcome dimension. Similarly, the reward field, *A*_x,*t*_, will be a vector, where each element of the vector denotes the amount of reward generated by earning a unit of the outcome along the corresponding dimension. As such, the total amount of reward earned by earning *δ***x** outcome will be *δ*x_1_[*A*_x,*t*_]_1_ + *δ*x_2_[*A*_x,*t*_]_2_ +…, which can be summarized as *δ***x**.*A*_x,*t*_. Similarly, the cost of earning outcomes will be a vector, where each element of the vector represents the cost of earning the corresponding outcome at the corresponding rate. Therefore the cost of earning *δ***x** amounts of outcome will be *δ***x**.*K*_v_. The net amount of reward will be:

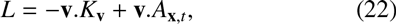

and the optimal trajectory will satisfy the Eular-Lagrange equation along each outcome dimension:

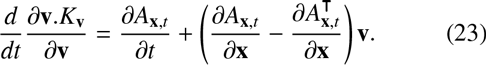

Furthermore since the end point of the trajectory is free (the total amount of outcomes is not fixed) we have:

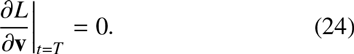

Here for simplicity we make the assumption that the cost of earning outcomes along each dimension is independent of the outcome rate on other dimensions, i.e.,

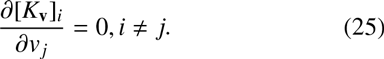

The above assumption implies that changing the rate of outcome along one outcome dimension will not affect the cost of earning outcomes along other dimensions. We will relax this assumption in the following sections. Under this assumption, the optimal speed will satisfy the following equation:

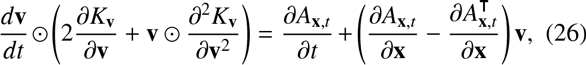

where ⊙ is the entrywise Hadamard product. See Appendix B for the proof.

To get a better intuition about the above equation and the optimal trajectory, here we present a geometrical interpretation. Let us assume that the cost function has the form indicated in equation 6, and also the time dependent changes in *A*_x,*t*_ are negligible (∂*A_x,t_*/∂*t* = 0). Under these conditions, if the outcome space is three dimensional, then equation 26 will be:

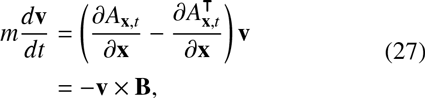

where × is the cross product, **B** is the curl of the reward field (**B** = `curl`*A*_x,*t*_), and *m* = 2*ak*^2^.^3^ Based on equation 27 the trajectory that the decision-maker takes in the outcome space can have two different forms, depending on whether **B** is zero. We will show in the next section that whenever the reward field is conservative (i.e., outcomes are substitutable for each other), then **B** will be zero and therefore the rate of earning outcomes will be constant throughout the session (*dv/dt* = 0). In contrast, in the condition that the reward field is non-conservative (i.e., outcomes are not substitutable for each other), **B** will be non-zero, and based on equation 27, the change in the rate of outcome earning will be perpendicular to the current rate of outcome earning. It can be shown that in this condition the decision-maker will take a circular trajectory in the outcome space (more details are provided in the next sections). Each of these two conditions are analyzed in turn in the next two sections.

### Conservative reward field

In line with the previous sections, let’s assume that *D*_x_ represents the motivational drive, and the amount of reward that consuming each outcome produces is equal to the amount of change that it makes in the motivational drive:

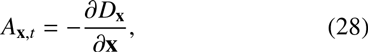

where *D*_x_ is a scalar field. We maintain the following theorem:

#### Theorem 3

*If the reward field is conservative, i.e., there exists a scalar field *D*_x_ such that equation 28 satisfies, and assuming that the time-dependent term of the reward field is zero* (∂*A_x,t_*/∂*t* = 0), *then the optimal rate of earning outcomes will be constant* (*d*v/*dt* = 0), *and it satisfies the following equation:*

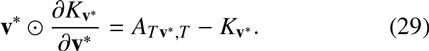

See Appendix B for the proof.

Therefore, if a reward field is conservative, then the optimal rate of earning outcomes is a constant rate. In the special case that the environment consists of two outcomes, the reward field being conservative implies that:

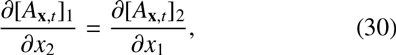

which suggests that the amount of decrement in the reward of the first outcome due to the consumption of the second outcome (∂[*A_x,t_*]_1_/∂*x_2_*), is equal to the decrement in the reward of the second outcome due to the consumption of the first outcome (∂[*A_x,t_*]_2_/∂*x_1_*); in other words, the reward field being conservative implies that the two outcomes have *the same degree of substitutability for each other*.

As an example, assume that the outcome space is two dimensional, and both outcomes satisfy the same internal state, say hunger. Consuming one unit of the first outcome decreases hunger by one unit, but consuming one unit of the second outcome reduces hunger by *l* units. As such, the motivationa*l* drive will be as follows:

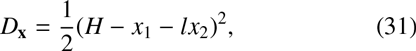

where *H* is the homeostatic set-point of hunger. Using equation 28 we have:

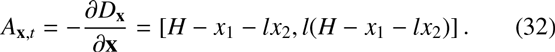

Based on Theorem 3, **B** will be zero (∂*A*_1_.∂*x*_2_ = ∂*A*_2_/∂*x*_1_) implying that the outcomes have substitutability for each other, and the response rate along each outcome dimension will be:

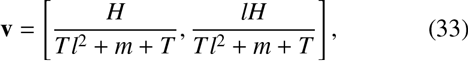

where *m* = 2*ak*^2^ and for simplicity it is assumed that *b* = 0. The above relation shows that the rate of earning each outcome is proportional to its rewarding effect, and inversely related to the cost of earning the outcome. Similar results can be obtained in the condition that the reward properties of the outcomes are the same, but the costs of earning the outcomes are different. For example, under a concurrent FR schedule in which an anima*l* needs to make *k* responses on one of the levers to earn outcomes, and *lk* responses on the other lever to earn the same outcome, the optimal rate of earning outcomes will be as follows:

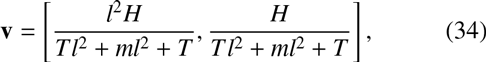

and the rate for each response will be:

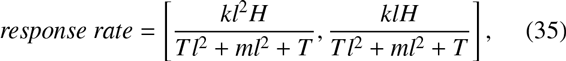

which states that the rate of responding for the first outcome (the outcome with the lower ratio-requirement) is *l* times greater than the second outcome. These results are generally in line with the probability matching notion, which states that a decision-maker allocates responses to outcomes based on the ratio of responses required for each outcome (Estes, 1950). Probability matching is generally considered as a violation of the rational choice theory, since within this theory it is expected that a decision-maker exclusively works for the outcome with the higher probability (the lower ratio requirement). However, based on the model proposed here, *probability matching* is the optimal strategy and therefore consistent with rational decision-making.

The above results, in fact, stem from the assumption that the cost of earning each outcome only depends on the rate of outcome earning in the same dimension, as expressed in equation 25. That is, the total cost of earning outcomes at rates *v*_1_ and *v*_2_ will be 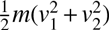, which implies that changing the response rate for one of the outcomes will not affect the cost of earning the other outcomes. As an example, imagine a concurrent instrumental conditioning experiment in which there are two levers available (left lever and right lever) and each lever leads to a different outcome. Here, the independence assumption entails that the cost of the current right lever press is determined by the time elapsed since the last right lever press and it does not matter whether there was a left lever press in between. Alternatively, one can assume what determines the cost is the delay between subsequent responses, either for the same or for a different outcome, i.e., the cost is proportional to the rate of earning all of the outcomes, and therefore, the cost takes the form 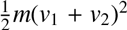. This alternative captures the fact that the cost is a function of the delay between subsequent responses, either for the same or for a different outcome. In this condition equation 25 does not hold anymore and the cost of earning outcomes will not be independent. Such a cost function can be achieved by defining the cost as follows:

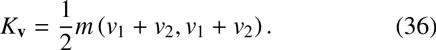

Given the above cost function, the optimal strategy is *maximisation*, i.e., to take the action with the higher reward (lower ratio-requirement), and to stop taking the other action:

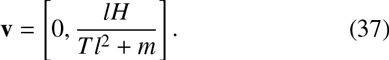

See Conservative reward field for the proof.

Therefore, whether the rate of outcome earning reflects probability matching or maximization strategy, depends on the cost function, and seemingly in the instrumental conditioning experiments, the cost that reflects the maximization strategy is better applicable. Regarding the experimental evidence, within the concurrent instrumental conditioning experiments, evidence in pigeons tested under the VR schedule (Herrnstein & Loveland, 1975) is in-line with the maximization strategy, and consistent with the prediction of the model. Within a wider scope of the decision-making tasks, some studies are consistent with the probability matching notion (Grant, Hake, & Hornseth, 1951) (see Vulkan (2000) for a review), while other studies provide evidence in-line with the maximization strategy (Edwards, 1961; Myers, Reilly, & Taub, 1961; Shanks, Tunney, & McCarthy, 2002; Siegel & Goldstein, 1959). Here, in most of the experiments the decision-making task used involves making a single choice (e.g., single button press) and receiving the feedback immediately (about whether the choice is rewarded), and after that the next trial starts. Such disjoint actions are unlikely to convey a rate-dependent cost, and, therefore, the structure of such studies do not readily fit in the model proposed here.

### Non-conservative reward field

In this section we assume that the reward field is nonconservative, i.e., there does not exist a scalar field *D*_x_ such that *A*_x,*t*_ satisfies equation 28. An example will be when the amount of reward that consuming an outcome produces is greater or smaller than the change in the motivational drive. For example, assume that there are two outcomes available, and the consumption of both outcomes has a similar effect on the motivational drive:

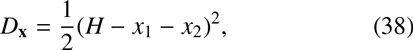

but the reward that the second outcome generates is *l* times larger than the change it creates in the motivational drive:

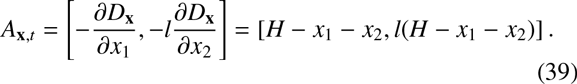

In this condition, ∂[*A*_x,*t*_]_1_/∂*x*_2_ = 1 and ∂[*A*_x,*t*_]_2_/∂*x*_1_ = *l*, and therefore the reward of the second outcome due to the consumption of the first outcome decreases more sharply than the reward of the first outcome would, due to the consumption of the second outcome. The reward field will be nonconservative and we have **B** = curl*A*_x,*t*_ = (0, 0, 1 – *l*), which will make the decision-maker take a circular trajectory in the outcome space. The exact characteristics of the optimal trajectory depend on the session duration and factor *l*, which is stated in the following theorem:

#### Theorem 4

*If the reward field follows equation 39 and the cost is as follows:*

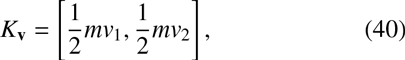

*then the optimal trajectory in the outcome space will be:*

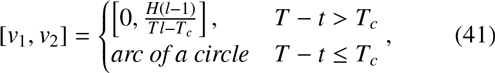

*where*

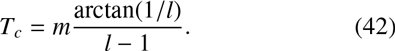

See Appendix B for the proof and more details about the optimal trajectory.

The above theorem implies that when the duration of the session is short (*T* < *T_c_*), then the trajectory in the outcome space is composed of a single segment, which is an arc of a circle. The response rate for both outcomes will be non-zero, and initially *v*_2_ > *lv*_1_, but at the end of the session the rate of earning the outcomes will be proportional to their reward effects (*v*_2_ = *lv*_1_), as indicated by equation 35 (Figure 4a,b). When the session duration is long, the trajectory in the outcome space will be composed of two segments. In the initial segment, the response rate for the outcome with the lower reward effect will be zero, and the decision-maker only works for the outcome with the higher reward effect. This segment continues until the time remaining to the end of the session is less than *T_c_*. After this time the second segment starts, which is an arc of a circle (Figure 4c).

A test of the prediction of Theorem 4, would be an experiment with two outcomes corresponding to the same food (and therefore having the same impact on the motivational drive) but with different levels of the desirability (e.g., two different flavors), and, therefore, having a different reward effect. We were not able to find such an experiment in the instrumental conditioning literature and, therefore, the prediction of Theorem 4 Theorem 4 will be left for future testing.

## Discussion

We formulated the problem of finding the optimal choice and response vigor in an optimal control framework by introducing the novel concept of reward field and using variational calculus methods. This formulation allowed us to derive the analytical solutions of the optimal rate of outcome earning in a wide range of experimental conditions. The analysis was divided into two sections: (1) the situations in which the environment contains only one outcome, and (2) the situations in which multiple outcomes can be earned in the environment. In the first condition, the results indicate that if the session duration is deterministic and known by the decision-maker, then the optimal rate of outcome earning is a constant rate throughout the session. Although these results are consistent with the majority of empirical results, they are inconsistent with the studies showing that the response rates decrease throughout an experiment session. We further showed that the uncertainty of the decision-maker about the session duration can explain this effect. In the conditions that the environment contains multiple outcomes, the results indicate that the optimal trajectory in the outcome space can take two different forms: (1) if the outcomes in the environment are substitutable for each other (equivalent to the reward field being conservative) then the optimal trajectory is a straight line in the outcome space; we discussed that these results are consistent with both probability matching and maximization notions of decision-making; (2) if the outcomes in the environment are not substitutable for each other, then the optimal trajectory is a straight line followed by a circular trajectory. To our knowledge, these results are the first analytical solution to the problem of choice and response vigor in the condition that the values of the outcomes can vary in the decision-making session.

**Figure 4.**
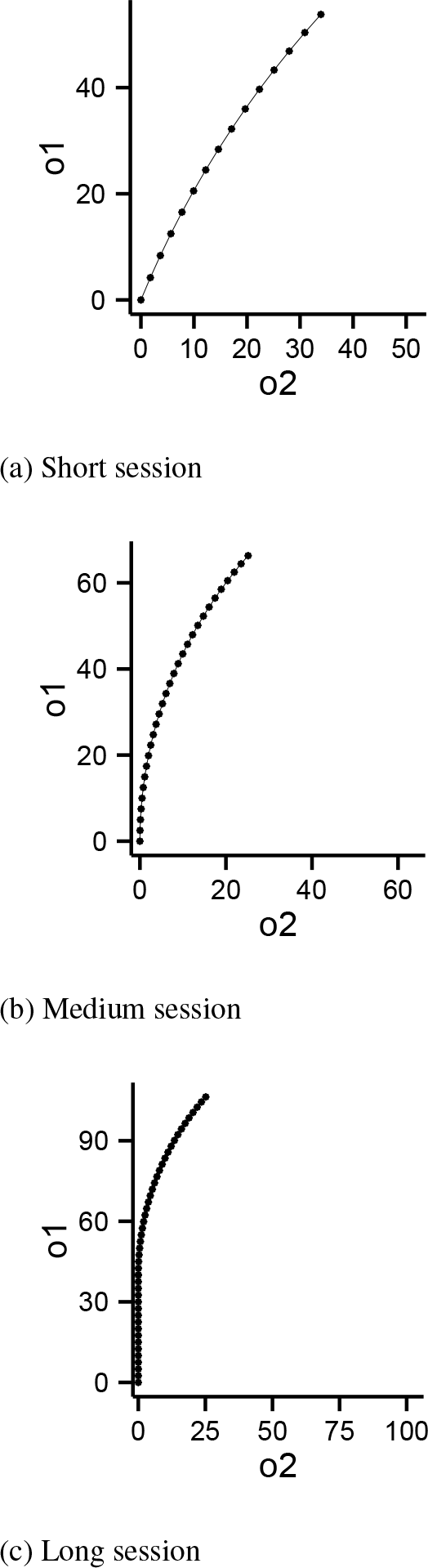
The optimal trajectories in the outcome space when the reward field is non-conservative. o1 and o2 are two different outcomes, where the amount of reward that *o*_2_ generates is larger than the decrement it creates in the motivational drive. Parameters used for simulations are *k* = 1, *l* = 1.1, *a* = 1, *b* = 0, *H* = 100, *m* = 2*ak*^2^ (a) Short session duration *T* = 7. (b) Medium session duration *T* = *T_c_* ≈ 14.75. (c) Long session duration *T* = 23.

There are two significant differences between the model proposed here and the previous models of response vigor (Dayan, 2012; Niv et al., 2007). Firstly, although the effect of between-session changes in the motivational drive on response vigor has been addressed in the previous models (Niv, Joel, & Dayan, 2006), the effects of the on-line changes in the motivational drive within a session are not addressed in the previous works, which we address in this model. Secondly, previous works conceptualized the structure of the task as a semi-Markov decision process, and derived the optimal actions that maximize the average reward per unit of time (average reward). Here, we used a variational analysis to calculate the optimal actions that maximize the reward earned within the session. One benefit of the approach taken in the previous works is that it extends naturally to a wide range of instrumental conditioning schedules such as interval schedules, while the extension of the model proposed here to the case of interval schedules is not trivial. Optimizing the average reward (as adopted in the previous works) is equivalent to the maximization of the reward in an infinite-horizon time scale, i.e., the session duration is unlimited; in contrast, the model used here explicitly represents the duration of the session, which as we showed plays an important role in the pattern of responses.

We assumed that the cost is only a function of the rate of earning outcomes, and it is time-independent (∂*k*_v_/∂_*t*_ = 0). However, in general one can assume that as time passes within a session, the cost of taking actions will be increased because of factors such as effector fatigue. Here we made the time-independence assumption based on previous studies that showed factors such as effector fatigue have a negligible effect on response rates (McSweeney, Hinson, & Cannon, 1996). Similarly, in the derivation of Theorem 2–4 we assumed that the time-dependent component of the reward field is zero (∂*A_x,t_*/∂*t* = 0), which seems to be a reasonable assumption, given the typical duration of an instrumental conditioning experiment.

The value of the outcomes can change because of factors other than changes in the motivational drive, such as specific satiety. In fact, the definition of the reward field is general and as long as it satisfies equations 1 and 2, the results obtained will be valid, irrespective of whether the underlying reason in the variability of the reward field is the motivational drive or other factors. Here, the reason that we used the motivational drive as the underlying cause of changes in the outcome values was because of the existence of the previous studies that link the quantity of outcome consumption to the reward of outcomes (Keramati & Gutkin, 2014); however, the model is general and can be applied to any other source that can cause changes in the value of the outcomes.

## Disclosures and Acknowledgments

NICTA is funded by the Australian Government as represented by the Dept. of Communications and the ARC through the ICT Centre of Excellence program. The funding source had no other role other than financial support.

All authors contributed in a significant way to the manuscript and that all authors have read and approved the final manuscript.

The author declares that the research was conducted in the absence of any commercial or financial relationships that could be construed as a potential conflict of interest.

## Appendix A Optimal trajectory in a single dimensional outcome space

**Derivation of cost earning outcomes**. The aim is to derive equation 6 from equation 5 under the assumption that *k* responses are required to earn one unit of the outcome.

*K_v_* is the cost of earning one unit of outcome at rate *v*. Earning the outcome at rate *v* implies that the time it takes to earn the outcome is 1/*v*, and since *k* responses have been executed in this period, the delay between responses is:

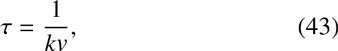

and therefore using equation 5, the cost of making one response will be *akv* + *b*. Since *k* responses are required for earning one unit of the outcome, the total cost will be *k* times the cost of one response:

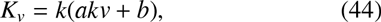

which is equivalent to equation 6.

In the case of non-deterministic schedules such as VR and RR schedules of reinforcement, the effect of actions on the rate of outcome earning will be non-deterministic, and therefore, the net reward (*L*) should be defined based on the expected rewards and costs:

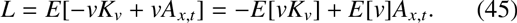

In the main text and in the following sections we use *v* instead of *E*[*v*] for simplicity of notation. Given this, the aim is to show that equation 45 is equivalent to equation 8. The second term in equation 45, i.e., *E*[*v*]*A*_*x,t*_, will be equivalent to *vA_x,t_* mentioned in equation 8. For the first term, i.e., *E*[*vK*_*v*_], we maintain that:

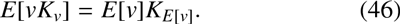

To show the above relation, let’s assume the subject performs *N* responses, and receives *r* outcomes. Since according to the definition of RR and VR schedules, out of *N* responses on average *N*=*k* will be rewarded, we have *E*[*r*] = *N/k* and the expected rate of outcome earning will be:

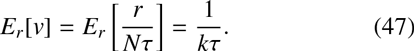

Therefore:

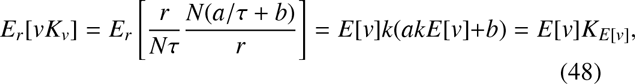

which proves equation 46.

**Fixed session duration**. The aim is to derive equation 11 and also to provide a proof for theorem 1. By substituting equation 8 in equation 10 we will have:

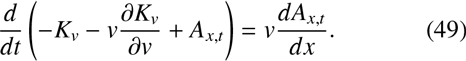

The term *dA_x,t_*/*dt* has two components: the first component is the change in *A_x,t_* due to the change in *x* and the second component is due to the time-dependent changes in *A_x,t_*:

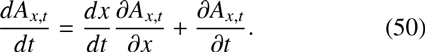

Furthermore we have:

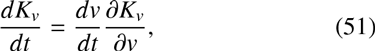

and similarly:

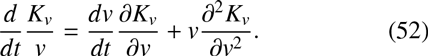

Substituting equations 50,51 and 52 in equation 49 yields:

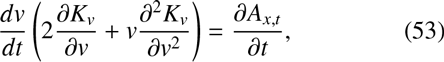

which is equivalent to equation 11. Given equations 2, 7 and 11 we will have:

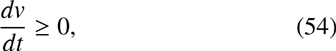

which proves the first part of Theorem 1. Assuming that ∂*A_x,t_*/∂*t* = 0, we are interested to find solutions to equation 11 and 12. One of the solutions to equation 11 is *dv/dt* = 0, and the other solution is:

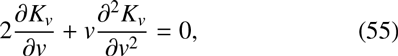

which is inconsistent with equation 7, and thus the only solution is *dv/dt* = 0. Given that the rate is constant we have *xT* = *vT*, which by substituting in yields equation 12, equation 13 which proves the second part of Theorem 1.

**Uncertain session duration**. Here we aim to derive the optimal trajectory when the duration of the session is probabilistic. The value function defined in equation 17 is as follows:

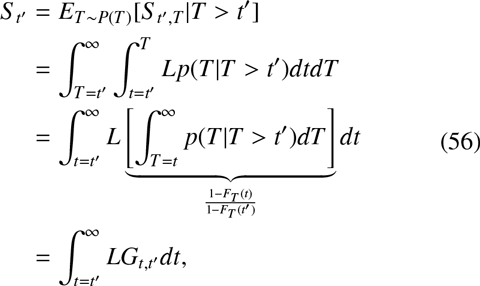

where *F_T_* is the cumulative probability distribution of *T*, and

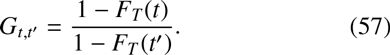

Using the Eular-Lagrange equation, the stationary solutionfor *S*’ will be:

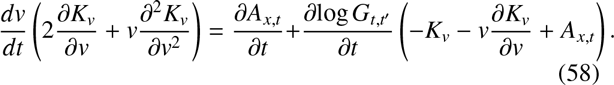

In the next section we provide an alternative method for approximating the optimal actions under uncertain session duration.

Expectation of session duration. The aim is to prove Theorem 2, and also to provide the details of the simulations. Assuming that the total session duration (*T*) has the probability density function *f(T)* and that *f(T)* > 0, here we show that the expectation of the session duration never decreases as time passes throughout the session. Let’s denote the expectation of the session duration at time *t*’ with *T*’:

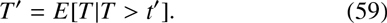

We have:

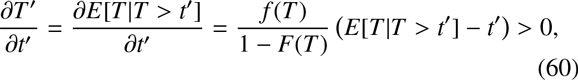

which implies that the expected duration of the session increases
as time passes.

Next, we show that the optimal response rate is a decreasing function of *t*’. Based on equation 12, the optimal response rate satisfies the following equation:

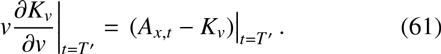

Taking the derivative w.r.t to *t*’ we get:

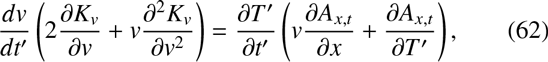

which given equations 1,60,7, and that *v* > 0, and assuming ∂*A_x,t_*/∂*T*′ yields:

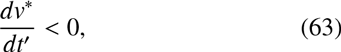

which implies that the rate of earning outcomes decreases as
time passes within a session.

For the simulation of the model, following McGuire and Kable (2013) we assumed that *T* follows a Generalized Pareto distribution:

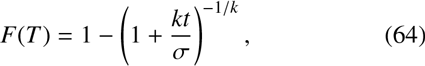

where *k* is a shape parameter and σ is a scale parameter, and the third parameter (location µ) was assumed to be zero. Furthermore we have:

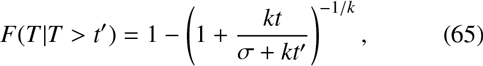

which has the following expected value:

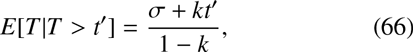

which as we expect is an increasing function of *t*’. For the simulation of the model we assumed that *k* = 0:9 and σ = 6, which represents that the initial expectation for the session duration is 60 minutes.

## Appendix B Optimal trajectory in a multi-dimension outcome space

The aim is to derive the optimal trajectory in the outcome space when there are multiple outcomes available (equation 23). The net reward at each point in time will be:

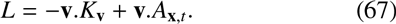

The Eular-Lagrange equation implies:

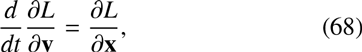

and therefore we have:

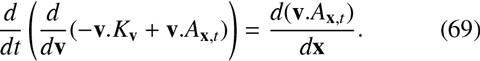

For the right hand side we have:

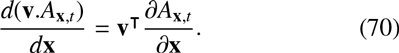

We also have:

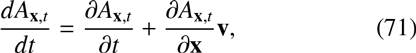

which by substitution into equation 69 we get:

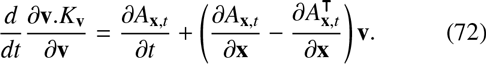

**Conservative reward field**. The aim is to prove Theorem 3, and also to derive equation 37. If the reward field is conservative, i.e., there exists a *D*_x_ such that equation 28 holds, we have:

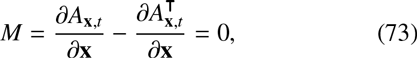

which is the consequence of the following:

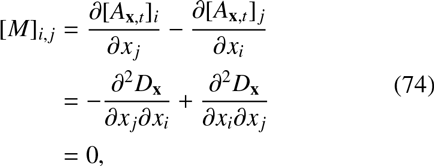

where we used equation 28 to derive the last equation. Based on equation 73, and assuming that, equation 72 will be:

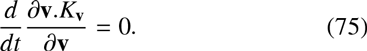

In the special condition that ∂*K*_v_/∂v is a diagonal matrix (i.e., equation 25 holds), equation 72 can be written as:

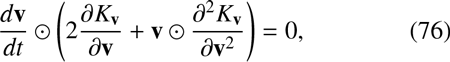

where ∂*K*_v_/∂v is a vector representing the diagonal terms of the matrix. For the derivation of the above equation we used the following relations:

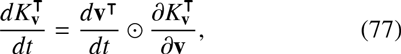

and:

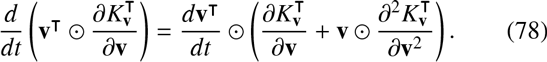

Given equation 7 the only admissible solution to equation 76 is *dv/dt* = 0, which shows that the optimal rate of earning outcomes is constant. By substituting equation 67 in the boundary conditions implied by equation 24 we get equation 29.

If ∂*K*_v_/∂v is not diagonal, and it is equivalent to the cost function shown in equation 36, then the Lagrangian (*L*) will be as follows:

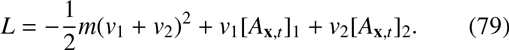

Using equation 75 we have:

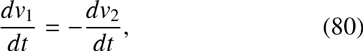

implying that

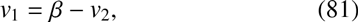

where β is a constant. Using boundary conditions in equation 24 we get at time *T*:

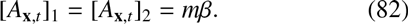

Given equation 32 and assuming that *l* ≠ 1 the only solution to the above equation is β = 0, which entails that *v*_1_ = −*v*_2_, which given the constrain that *v*_1_ ≥ 0 and *v*_2_ ≥ 0 is not an admissible solution. Assuming that v1 takes the boundary value v1 = 0, the problem degenerates into a problem involving only one outcome (since *v*_1_ = 0) which can be solved using Theorem 1. Solving equation 13, assuming *b* = 0 and *m* = 2*ak*^2^, yields equation 37.

**Non-conservative reward field**. The aim is to prove Theorem 4. We have:

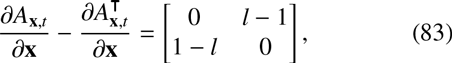

and based on equation 26 we get:

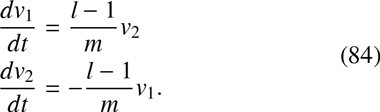

Defining *w* = (*l* − 1)/*m*, the solution to the above set of differential equations has the form:

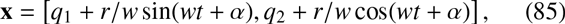

which is an arc of a circle centered at [*q*_1_; *q*_2_], and *r* and *α* are free parameters. The parameters can be determined using the boundary condition imposed by equation 24, and also assuming that the initial position is **x** = 0. The boundary condition in equation 24 implies:

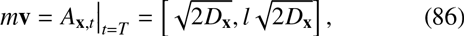

which implies that at the end of the trajectory the rate of earning the second outcome is *l* times larger than the first outcome. Therefore, the general from of the trajectory will be an arc starting from the origin and ending along the above direction. Given the constrain that *v* ≥ 0 only the solutions in which *q*_2_ ≤ 0 are acceptable ones (i.e., the center of the circle is below the x-axis). Solving equation 85 for *q*_2_ ≤ 0 we get:

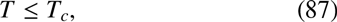

where

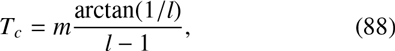

and therefore *T_c_* is independent of *H* (the initial motivational drive). As such if *T* satisfies equation 87 then the optimal trajectory will be an arc of a circle starting from the origin. Otherwise, if *T* > *T_c_*, the optimal trajectory will be composed of two segments. In the first segment, *v*_1_ will take the boundary condition *v*_1_ = 0 and the decision-maker earns only the second outcome (the outcome with the higher reward effect). The first segment continues until the remaining time in the session satisfies equation 87 (the remaining time is less than *T_c_*), after which the second segment starts, which is an arc of a circle defined by equation 85. The rate of earning the second outcome, *v*_2_, in the first segment of the trajectory (when *v*_1_ = 0) can be obtained by calculating the rates at the beginning of the circular segment. The initial rate at the start of the circular segment is as follows:

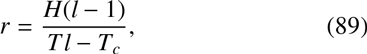

which implies that at the first segment of the trajectory we have:

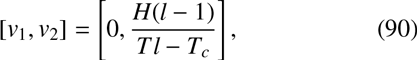

which completes the proof of Theorem 4.

## Footnotes

1 In equation 14 the optimal outcome rate is dependent on the initial drive (*H*) and the session duration (*T*), and therefore one might intuitively think that if for example half-way through the session the decision-maker re-calculates the optimal rate using equation 14, then it will get a different constant rate than the one it got initially at the beginning of the session, as *H* and *T* will be different from the ones at the start of the session. This argument, however, is not correct, and under the optimal rate, the motivational drive and the remaining time in the session change in opposite directions, in a way that the optimal rate remains invariant throughout the session, irrespective of what time in the session equation 14 is calculated.

2 Note that in the original notation in Killeen and Sitomer (2003), *α* is denoted by *a* and β is denoted by *b*.

3 It is interesting to note that equation 27 in fact lays out the motion of a unit charged particle (negatively charged) with mass m in a magnetic field with magnitude **B**.

## References

Aberman, J. E., & Salamone, J. D. (1999). Nucleus accumbens dopamine depletions make rats more sensitive to high ratio requirements but do not impair primary food reinforcement. Neuroscience, 92(2), 545–52.

Adair, E. R., & Wright, B. A. (1976). Behavioral thermoregulation in the squirrel monkey when response effort is varied. Journal of Comparative and Physiological Psychology, 90(2), 179.

Alling, K., & Poling, A. (1995, may). The effects of differing response-force requirements on fixed-ratio responding of rats. Journal of the experimental analysis of behavior, 63(3), 331–46.

Barofsky, I., & Hurwitz, D. (1968). Within ratio responding during fixed ratio performance. Psychonomic Science, 11(7), 263–264.

Baum, W. M. (1993). Performances on ratio and interval schedules of reinforcement: Data and theory. Journal of the Experimental Analysis of Behavior, 59(2), 245.

Dayan, P. (2012, apr). Instrumental vigour in punishment and reward. The European journal of neuroscience, 35(7), 115–268.

Edwards, W. (1961). Probability learning in 1000 trials. Journal of Experimental Psychology, 62(4), 385.

Estes, W. K. (1950). Toward a statistical theory of learning. Psychological review, 57(2), 94.

Felton, M., & Lyon, D. O. (1966). The post-reinforcement pause. Journal of the Experimental Analysis of Behavior, 9(2), 131–134.

Ferster, C. B., & Skinner, B. F. (1957). Schedules of reinforcement.Prentice-hall inc.

Foster, M., Blackman, K., & Temple, W. (1997). Open versus closed economies: performance of domestic hens under fixed ratio schedules. Journal of the Experimental Analysis of Behavior, 67(1), 67.

Grant, D. A., Hake, H. W., & Hornseth, J. P. (1951). Acquisition and extinction of a verbal conditioned response with differing percentages of reinforcement. Journal of experimental psychology, 42(1), 1.

Greenwood, M. R., Quartermain, D., Johnson, P. R., Cruce, J. A., & Hirsch, J. (1974, nov). Food motivated behavior in genetically obese and hypothalamic-hyperphagic rats and mice. Physiology & behavior, 13(5), 687–92.

Herrnstein, R. J., & Loveland, D. H. (1975). Maximizing and matching on concurrent ratio schedules. Journal of the experimental analysis of behavior, 24(1), 107.

Hull, C. L. (1943). Principles of Behavior. New York: Appleton-Century.

Keesey, R. E., & Kling, J. W. (1961). Amount of reinforcement and free-operant responding. Journal of the Experimental analysis of behavior, 4(2), 125–132.

Kelsey, J. E., & Allison, J. (1976). Fixed-ratio lever pressing by VMH rats: Work vs accessibility of sucrose reward. Physiology & behavior, 17(5), 749–754.

Keramati, M. (2011). A Reinforcement Learning Theory for Homeostatic Regulation‥

Keramati, M., & Gutkin, B. (2014). Homeostatic reinforcement learning for integrating reward collection and physiological stability. eLife, 3.

Killeen, P. R. (1994, feb). Mathematical principles of reinforcement. Behavioral and Brain Sciences, 17, 105–172.

Killeen, P. R. (1995a). Economics, ecologics, and mechanics: The dynamics of responding under conditions of varying motivation. Journal of the Experimental Analysis of Behavior, 64(3), 405–431.

Killeen, P. R. (1995b, nov). Economics, ecologics, and mechanics: The dynamics of responding under conditions of varying motivation. Journal of the Experimental Analysis of Behavior, 64(3), 405–431. 10.1901/jeab.1995.64-405

Killeen, P. R., & Sitomer, M. T. (2003, apr). MPR. Behavioural Processes, 62(1–3), 49–64.

Liberzon, D. (2011). Calculus of Variations and Optimal Control Theory: A Concise Introduction. Princeton University Press.

Lowe, C. F., Davey, G. C. L., & Harzem, P. (1974). Effects of reinforcement magnitude on interval and ratio schedules. Journal of the Experimental analysis of behavior, 22(3), 553–560.

Mazur, J. E. (1982). A molecular approach to ratio schedule performance. In M. L. Commons, R. J. Herrnstein, & H. Rachlin (Eds.), Quantitative analyses of behavior vol. 2: Matching and maximizing accounts. Ballinger.

McGuire, J. T., & Kable, J. W. (2013, apr). Rational temporal predictions can underlie apparent failures to delay gratification. Psychological review, 120(2), 395–410.

McSweeney, F. K. (2004). Dynamic changes in reinforcer effectiveness: Satiation and habituation have different implications for theory and practice. The Behavior Analyst, 27(2), 171–188.

McSweeney, F. K., & Hinson, J. M. (1992, jul). Patterns of responding within sessions. Journal of the Experimental Analysis of Behavior, 58(1), 19–36.

McSweeney, F. K., Hinson, J. M., & Cannon, C. B. (1996). Sensitization-habituation may occur during operant conditioning. Psychological Bulletin, 120(2), 256.

McSweeney, F. K., Roll, J. M., & Weatherly, J. N. (1994, jul). Within-session changes in responding during several simple schedules. Journal of the Experimental Analysis of Behavior, 62(1), 109–132.

McSweeney, F. K., Weatherly, J. N., & Swindell, S. (1995, jul). Within-session changes in key and lever pressing for water during several multiple variable-interval schedules. Journal of the Experimental Analysis of Behavior, 64(1), 75–94.

Meunier, G. F., & Starratt, C. (1979). On the magnitude of reinforcement and fixed-ratio behavior. Bulletin of the Psycho-nomic Society, 13(6), 355–356.

Myers, J. L., Reilly, R. E., & Taub, H. A. (1961). Differential cost, gain, and relative frequency of reward in a sequential choice situation. Journal of experimental Psychology, 62(4), 357.

Neumann, L. J., & Morgenstern, O. (1947). Theory of games and economic behavior (Vol. 60). Princeton university press Princeton.

Niv, Y., Daw, N. D., Joel, D., & Dayan, P. (2007, apr). Tonic dopamine: opportunity costs and the control of response vigor. Psychopharmacology, 191(3), 507–20.

Niv, Y., Joel, D., & Dayan, P. (2006, aug). A normative perspective on motivation. Trends in cognitive sciences, 10(8), 375–81. 10.1016/j.tics.2006.06.010

Pear, J. (2001). The science of learning. Psychology Press.

Powell, R. W. (1968). The effect of small sequential changes in fixed-ratio size upon the post-reinforcement pause. Journal of the Experimental Analysis of Behavior, 11(5), 589–593.

Powell, R. W. (1969). The effect of reinforcement magnitude upon responding under fixed-ratio schedules. Journal of the Experimental Analysis of Behavior, 12(4), 605–608.

Premack, D., Schaeffer, R. W., & Hundt, A. (1964). Reinforcement of drinking by running: effect of fixed ratio and reinforcement time. Journal of the experimental analysis of behavior, 7(1), 91–96.

Rachlin, H. (2000). The Science of Self-Control. Harvard University Press.

Salimpour, Y., & Shadmehr, R. (2014, jan). Motor costs and the coordination of the two arms. The Journal of neuroscience: the official journal of the Society for Neuroscience, 34(5), 1806–18.

Shanks, D. R., Tunney, R. J., & McCarthy, J. D. (2002). A re-examination of probability matching and rational choice. Journal of Behavioral Decision Making, 15(3), 233–250.

Sidman, M., & Stebbins, W. C. (1954). Satiation effects under fixed-ratio schedules of reinforcement. Journal of Comparative and Physiological Psychology, 47(2), 114.

Siegel, S., & Goldstein, D. A. (1959). Decision-making behavior in a two-choice uncertain outcome situation. Journal of Experimental Psychology, 57(1), 37.

Vulkan, N. (2000). An economist’s perspective on probability matching. Journal of economic surveys, 14(1), 101–118.

